# A universal buffer system for native LC-MS analysis of antibody-based therapeutics

**DOI:** 10.1101/2025.11.17.688837

**Authors:** Kent R. Vosper, Bradley T. V. Davis, Jagandeep Saraya, Derek K. O′Flaherty, Algirdas Velyvis, Siavash Vahidi

## Abstract

Liquid chromatography coupled to mass spectrometry (LC-MS) is a powerful analytical technique for analyzing biological macromolecules. A long-standing challenge has been applying LC-MS at physiological pH under native conditions using volatile buffers. The predominant “buffer” used, ammonium acetate (AmAc, p*K*_a_ 4.75 for acetic acid and 9.25 for ammonium), does not offer sufficient buffering capacity in the physiological pH range of 7.0–7.4. To address this, we evaluated a set of fluorinated ethylamines, 2-fluoroethylamine (MFEA, p*K*_a_ 8.9), 2,2-difluoroethylamine (DFEA, p*K*_a_ 7.2), and 2,2,2-trifluoroethylamine (TFEA, p*K*_a_ 5.5), that together provide buffering across the 4.5-9.8 pH range. We show that protein separations on strong cation- and anion-exchange resins in these volatile mobile phases perform comparably to traditional non-volatile buffers, with similar elution profiles and analyte elution ranking, albeit with slightly broader peaks. Using fully volatile gradients of pH or ionic strength, we chromatographically resolved charge variants of protein analytes such as mAbs and bovine serum albumin. For many of the eluting LC peaks, we obtained high-resolution mass spectra capable of resolving glycoforms of antibodies. Hydrophobic interaction chromatography (HIC) in volatile mobile phases preserved native separation order and further resolved drug-to-antibody ratio (DAR) species of the antibody-drug conjugate brentuximab-vedotin. For each chromatography modality we further compare innovator and biosimilar antibodies, demonstrating the reproducibility of results in the proposed volatile compounds. Together, our results establish fluorinated ethylamines, in combination with ammonium acetate, as a universal volatile buffer system for native LC-MS, broadly applicable across major chromatographic modalities while maintaining compatibility with mass spectrometry.

## Introduction

Monoclonal antibody (mAb) therapeutics have emerged as highly effective treatments for cancer, autoimmune diseases, and other conditions because of their ability to bind specific cell-surface antigens with precision. Examples include trastuzumab, which targets HER2;^1^ bevacizumab, which blocks VEGF-A–driven angiogenesis;^2^ and pembrolizumab, which inhibits an immune checkpoint.^3^ Other antibodies act directly in circulation, such as eculizumab, which binds complement component C5.^4^ Antibody–drug conjugates (ADCs) extend this principle further by linking antibodies to cytotoxic payloads, as in brentuximab-vedotin, which delivers monomethylauristatin A to CD30-positive tumor cells.^5–7^ The therapeutic utility of mAbs and ADCs underscores the importance of rigorous quality control, since aggregation and structural heterogeneity can compromise efficacy and safety.^8–11^ Liquid chromatography (LC) has become a cornerstone of this process, enabling rapid and versatile analysis of multiple protein properties.^8–11^

Coupling LC to mass spectrometry (LC-MS) adds a powerful orthogonal readout by combining separation based on analyte physicochemical properties with direct measurement of molecular mass. Reverse-phase (RP) LC-MS^12^ is well established for intact mass analysis of mAbs proteoforms, but it does not report on native-state properties such as oligomeric state, conformation, complex formation, or aggregation as it is often performed under denaturing conditions using an acetonitrile gradient, and typically in the presence of formic or trifluoroacetic acid.^13^ Native LC-MS seeks to recover this information by pairing electrospray ionization (ESI) with chromatographic modes such as size-exclusion (SEC), ion exchange (IEX), and hydrophobic interaction chromatography (HIC).^14–21^ These modalities are indispensable for antibody characterization. For instance, HIC provides critical resolution of drug-to-antibody ratio (DAR) species during ADC development.^21–23^ However, use of these LC modalities in LC-MS is hindered by non-volatile salts and buffers, which cause severe adduction and peak broadening in ESI spectra.

The scarcity of MS-compatible, volatile buffering compounds has led to the widespread adoption of ammonium acetate (AmAc) as the most common “buffering” component when coupling protein LC to MS.^24–27^ Despite its utility, AmAc provides little buffering capacity in the physiological range of pH 7.0–7.4,^28,29^ given the p*K*_a_ values of its conjugate acid/base pairs (4.75 for acetic acid and 9.25 for ammonium). This gap is problematic because ESI is prone to electrochemically-driven pH shifts of up to four units,^30,31^ which can destabilize proteins and introduce artefacts into the data. To address this, we recently introduced two fluorinated ethylamines, 2,2-difluoroethylamine (DFEA, p*K*_a_ 7.2) and 2,2,2-trifluoroethylamine (TFEA, p*K*_a_ 5.5), as volatile pH buffers compatible with native MS.^31^ Here, we expand this system by introducing a third compound, 2-fluoroethylamine (MFEA, predicted p*K*_a_ 9.0)^32^, alongside DFEA and TFEA, in LC-MS workflows. We demonstrate their compatibility with SEC, IEX, and HIC, thereby establishing a universal volatile buffer system that extends the utility of native LC-MS beyond the limitations of AmAc alone.

## Results

### Cation Exchange Chromatography (CEX) with an ionic strength gradient

Because IEX depends critically on precise control of analyte ionization, we reasoned that it would directly benefit from the enhanced buffering capacity provided by the fluorinated ethylamines. To test this, we developed a workflow in which each condition was evaluated under both a steep gradient (1 M salt over 14 minutes) and a shallow gradient for higher resolution (200 mM salt over 20 or 30 minutes). This experimental design allowed us to directly compare chromatographic performance under conditions emphasizing either speed or resolution. We also assessed whether pH gradients could be used in place of traditional ionic strength gradients, employing volatile buffers set at pH 5.46 and 9.84. This approach provided a direct test of the buffering range afforded by the fluorinated ethylamines in an IEX setting.

We began by validating previously reported results^33^ for mAb CEX using a MES-based pH 6 buffer. These conditions produced strong binding of basic mAbs (trastuzumab, NISTmAb, and bevacizumab), as well as cytochrome *c*, to a Waters Protein-Pak Hi Res SP resin (Figure 1A and Figure S1A). When elution was performed with a shallow 0-200 mM gradient, we observed the expected order of elution: trastuzumab, NISTmAb, and followed by bevacizumab, consistent with the literature.^34–37^ Although bevacizumab (pI 8.3) has a lower reported pI than trastuzumab (pI 9.1) and NISTmAb (pI 9.2), it consistently eluted after NISTmAb, a trend noted previously.^38^ Cytochrome *c* (pI 9.5) required a higher ionic strength (100-300 mM) for elution, consistent with its strongly basic nature.

**Figure 1.**
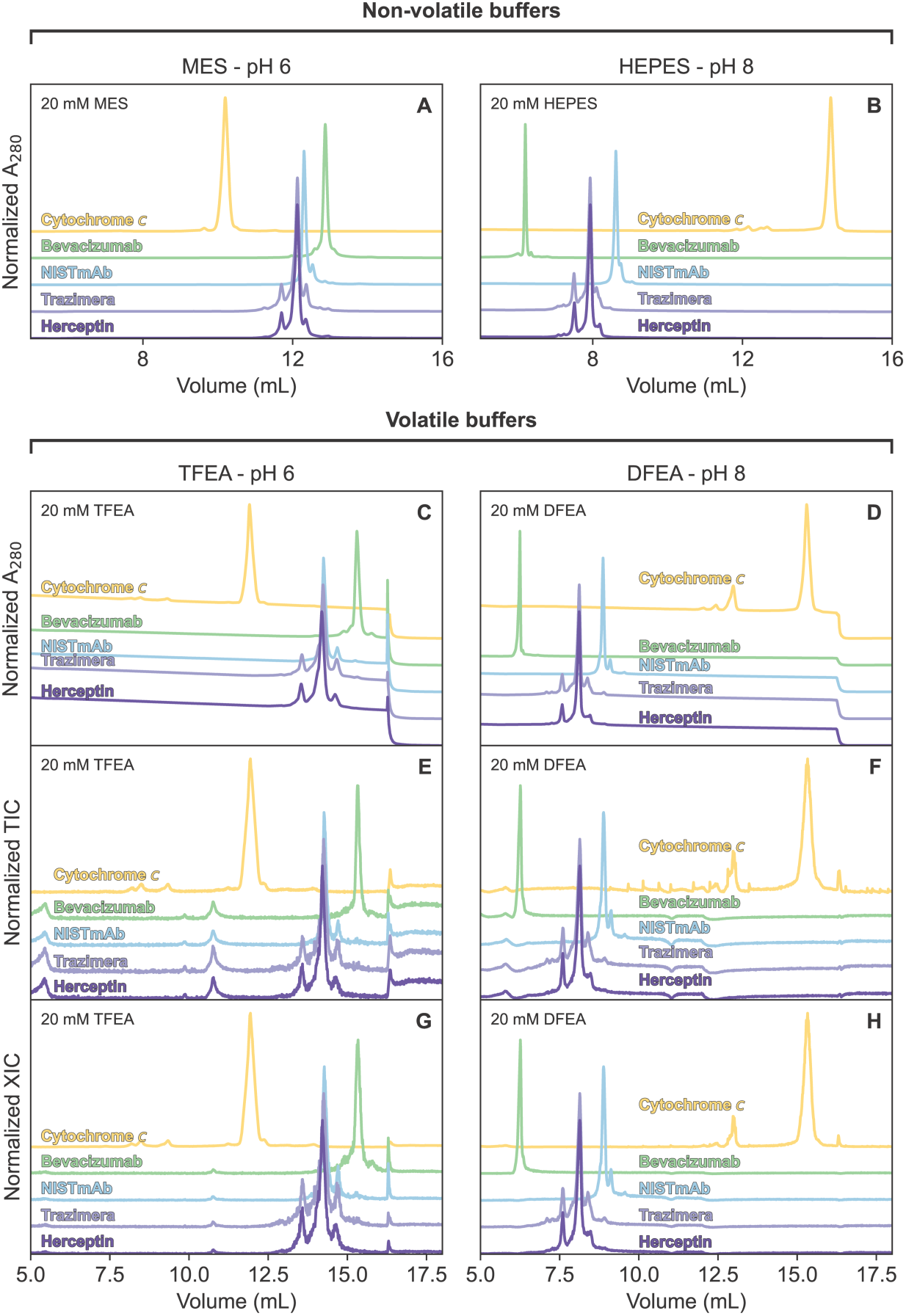
Cation exchange chromatography at pH 6 and 8 comparing TFEA and DFEA to non-volatile buffering compounds. Traces are colour coded for each analyte. A_280_ chromatograms obtained with an MES mobile phase (pH 6.0) (A) or a HEPES mobile phase (pH 8.0) (B), and a shallow NaCl gradient. The next rows of plots show A_280_ (C, D), TIC (E, F), and XIC (G, H) obtained with either a TFEA mobile phase (pH 6.0) or a DFEA mobile phase (pH 8.0), also using a shallow ionic strength gradient produced with AmAc. Baselines for each analyte have been offset to improve readability of overlapping peaks.

To directly compare the proposed volatile buffers with traditional non-volatile systems, we replaced MES with TFEA and NaCl with AmAc, thereby maintaining ionic strength while ensuring ESI-MS compatibility. The resulting chromatograms closely mirrored those obtained with MES/NaCl, with only minor shifts in retention time (Figure 1C). Notably, small secondary peaks arising from acidic and basic variants of the mAbs were resolved just as effectively in the volatile mobile phase, demonstrating that TFEA/AmAc can faithfully reproduce established CEX behavior at pH 6.0.^38,39^ We extended these experiments to pH 8.0, substituting HEPES with DFEA in the mobile phase and again testing both steep and shallow salt gradients. At this higher pH, absorbance chromatograms measured at 280 nm aligned well between volatile and non-volatile phases (Figure 1D). However, in the LC-MS total ion chromatograms (TICs), additional non-proteinaceous peaks were present (Figure 1E and F). Because these peaks did not absorb at 280 nm and were consistently present across injections, we attribute them to low-level contaminants in the reagent-grade fluorinated ethylamines (MS-grade reagents are not currently available). Importantly, plotting extracted ion chromatograms (XICs) of the protein signals eliminated these spurious features (Figure 1G and H). To test reproducibility, we analyzed trastuzumab from two different suppliers (Herceptin versus the biosimilar Trazimera) and observed nearly identical peak shapes, heights, and retention times using our volatile buffer conditions. When analyzing cytochrome *c* at pH 8, we noted the presence of a small early-eluting peak, already detectable in HEPES buffer but more pronounced in DFEA. This effect may reflect subtle differences in how DFEA influences protein structure or its interactions with the resin, but the main chromatographic features were well preserved. These conclusions also held for our steep gradients with a modest loss of chromatographic resolution (Figure S1).

To evaluate the compatibility of the proposed fluorinated ethylamine buffers with native ESI-MS, we examined the resulting mass spectra. We integrated the entire width of the main LC peak for each analyte and inspected the raw and deconvoluted mass spectra recorded using TFEA (pH 6) and DFEA (pH 8). We observed excellent signal quality and resolution for cytochrome *c* (Figure 2A and B) and across all tested antibody-based analytes (Figure 2C-H). Different patterns of glycosylation variants are apparent, spaced by approximately 162 Da in deconvoluted spectra of our tested antibodies. Four main mass peaks (148034, 148196, 148358, and 148518 Da) of NISTmAb observed in this CEX correspond closely to the masses we measured by RP-LC-MS analysis (148038, 148199, 148361, 148522 Da) of the same sample (Figure 2E and F), which have been assigned to glycoforms G0F/G0F, G0F/G1F, G1F/G1F, G1F/G2F in the literature.^40^ While it is well resolved in LC dimension, the peak of the basic variant of NISTmAb is lower intensity, therefore the resulting mass spectrum is rather noisy resulting in larger uncertainty in deconvoluted mass (Figure S2). Nevertheless, the mass difference of approximately 130 Da between main and minor LC proteoforms is apparent, most likely corresponding to an addition of a single C-terminal lysine residue, a well-known mAb modification. The S/N ratio in the mass spectra of minor charge variants of trastuzumab was not sufficient to obtain accurate mass measurements for these species (Figure 2G and H).

**Figure 2:**
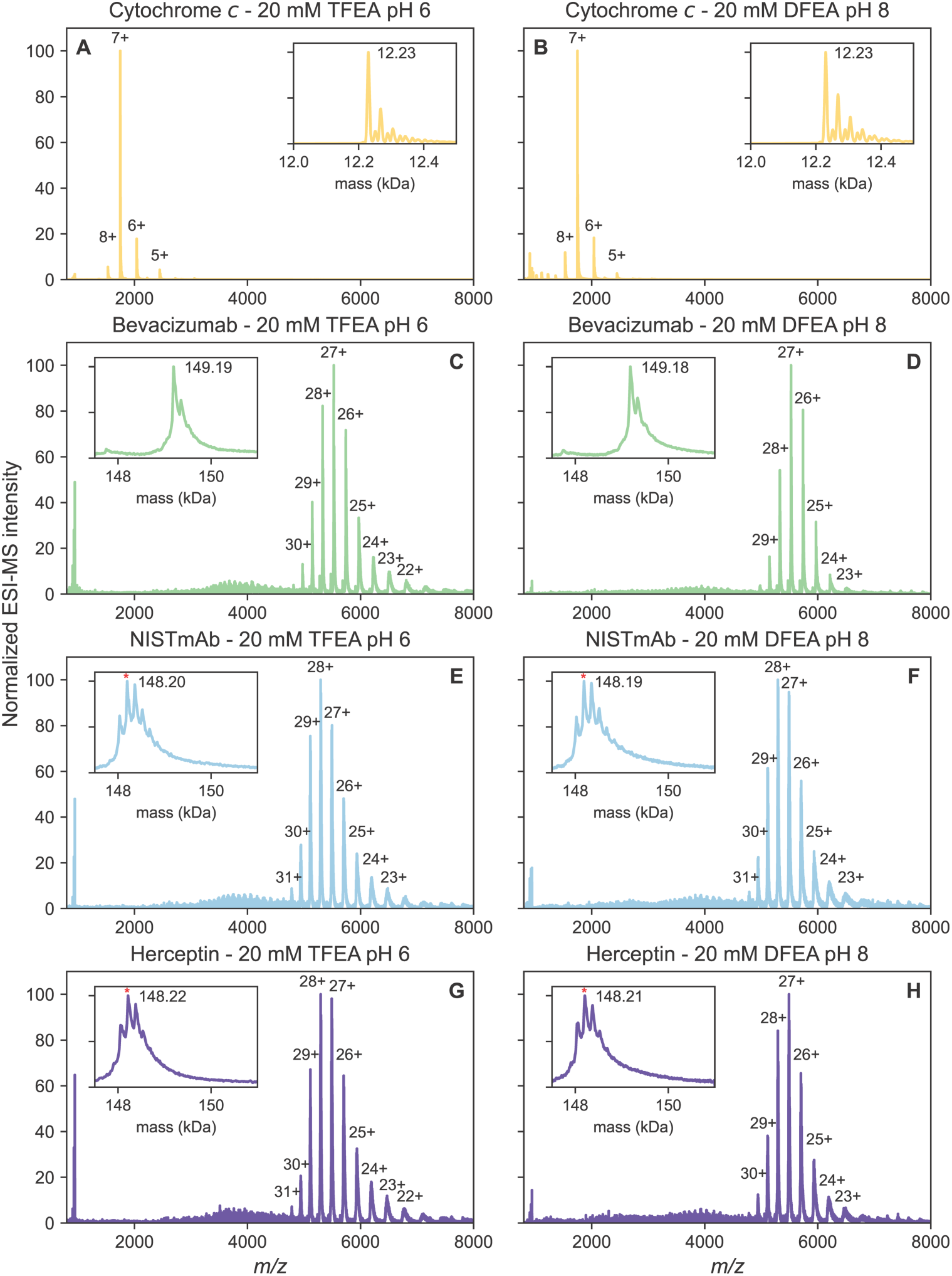
Mass spectra recorded during pH 6 and pH 8 CEX chromatography MS. Spectra and deconvoluted mass plots colour coded for each analyte. Plots are displayed for spectra obtained in both pH 6 (A, C, E, G) and pH 8 (B, D, F, H). Prominent charge states are labelled and deconvoluted masses are displayed as inset plots. In deconvolution plots for Herceptin and NISTmAb the highest mass peak is denoted with a red asterisk.

### Anion Exchange Chromatography (AEX) with an ionic strength gradient

We first validated the buffering properties of 2-fluoroethylamine (MFEA; we prefer to include prefix “mono-” in the abbreviated name as it allows to refer to each of the fluorinated ethylamines without ambiguity), which has been reported in the medicinal chemistry literature with a *pK*_a_ of 9.0, though without experimental details.^32^ We performed a potentiometric pH titration and fit the resulting curve to a single-*pK*_a_ model, yielding a value of 8.93 ± 0.1 for the conjugate acid of MFEA (Figure S3). By the standard rule that effective buffering occurs within one pH unit of the *pK*_a_, MFEA can provide buffering in the range of ∼8-10. Together with DFEA (*pK*_a_ 7.2), these compounds therefore extend volatile buffering capacity across the neutral-to-basic window of pH 6.2-9.9, a range often critical for biochemical studies.

We next verified the performance of MFEA in AEX chromatography using a Waters Protein-Pak Hi Res Q column. At pH 9, MFEA-based buffers were compared against traditional Tris buffers with a shallow salt gradient using NaCl for the non-volatile control and AmAc for the volatile mobile phases. As in the CEX experiments, retention times were slightly shifted under conditions containing volatiles, but the elution order was preserved and corresponded well to the pI value of the analytes: NISTmAb (very basic, minimal retention), trastuzumab (innovator and biosimilar co-eluting), bevacizumab (less basic), and acidic eculizumab, which required higher salt concentrations for elution. We also observed strong retention for bovine serum albumin (BSA) and the antibody-drug conjugate brentuximab-vedotin, which uniquely showed slightly stronger binding in volatile mobile phases compared to Tris-NaCl system (Figure 3). Importantly, high-quality mass spectra were obtained for all mAb analytes under volatile conditions, confirming the MS compatibility of MFEA (Figure 4 and Figure S4). These conclusions also held for our steep gradients with a modest loss of chromatographic resolution (Figure S5). Brentuximab-vedotin, an antibody-drug conjugate, was a uniquely challenging analyte in AEX analysis, since the protein spectrum is split into multiple components due to heterogeneity in the number of attached drug molecules, as well as differential glycosylation. Nevertheless, we were able to record a mass spectrum for this mixture of molecular species (Figure 4C). Despite the fact that the deconvoluted spectrum is affected by increased noise level, the three main peaks at 150880, 153510, and 156160 Da correspond well to the masses expected for mAb with 2, 4, and 6 monomethylauristatin A molecules (MW = 1316 Da) attached. Moreover, each of the main peaks shows a fine structure of signals differing by ∼160 Da, corresponding to a mixture of G0F/G1F glycosylated species (Figure S4D). Drug-to-antibody ratio was investigated even more effectively, and AEX mass measurement were independently validated, by the hydrophobic interaction chromatography (see below).

**Figure 3.**
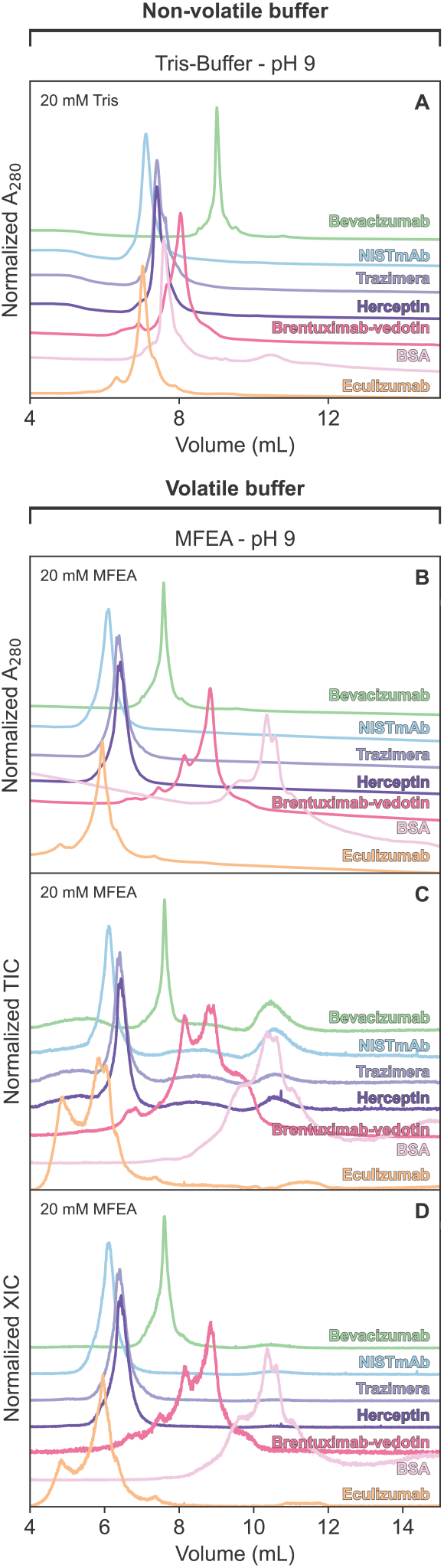
Anion exchange chromatography at pH 9 comparing MFEA to non-volatile TrisHCl-based buffer. Traces are colour coded for each of the analytes. The first plot displays an A_280_ chromatogram (A) obtained in a non-volatile mobile phase, Tris at pH 9, with a shallow ionic strength gradient. The A_280_ (B), TIC (C), and XIC (D) data obtained in MFEA buffer at pH 9 are shown in the subsequent plots. Baselines for each analyte have been offset to improve readability of overlapping peaks.

**Figure 4:**
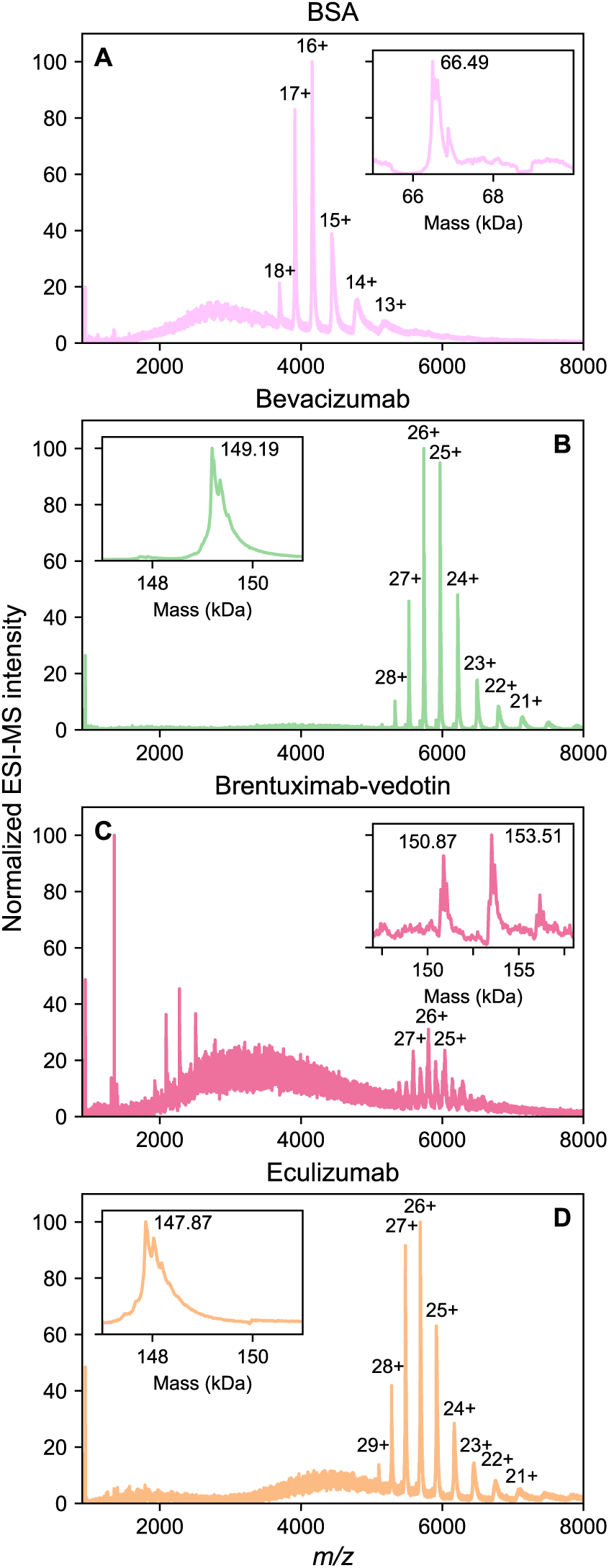
Mass spectra recorded during pH 9 AEX chromatography MS. Spectra and deconvoluted mass traces are colour coded for each analyte. Prominent charge states are labelled and deconvoluted masses are displayed as inset plots.

Together, these results establish MFEA as an effective volatile buffering component for AEX chromatography, enabling seamless integration with MS without compromising chromatographic resolution or spectral quality.

### Ion Exchange Chromatography (IEX) with a pH gradient

The series of ethylamine and its fluorinated derivatives (ethylamine, p*K*_a_ 10.7; MFEA, pK_a_ 8.9; DFEA, p*K*_a_ 7.2; TFEA, p*K*_a_ 5.5) provides a set of weak bases whose conjugate acids span a wide p*K*_a_ range, enabling precise formulation of mobile phases across pH 5-10. When combined with acetic acid (p*K*_a_ 4.75) or formic acid (p*K*_a_ 3.75) for titration, the resulting mixtures are fully volatile and allow finely controlled pH gradients that remain compatible with MS detection. To evaluate this approach, we prepared volatile buffer systems representing low pH (40 mM AcOH, 20 mM TFEA, 20 mM DFEA, pH 5.46) and high pH (32 mM AcOH, 20 mM DFEA, 20 mM MFEA, 30 mM ethylamine, pH 9.84) conditions. When applied to CEX chromatography using a Waters Protein-Pak Hi Res SP column, sweeping a gradient from pH 5.46 to 9.84 resulted in a chromatogram where each UV A_280_ signal corresponded to a TIC peak (Figure 5A and B). For brentuximab-vedotin, the TIC was of lower quality, consistent with baseline issues also visible in the A_280_ trace; however, applying a mass filter recovered a clean extracted ion chromatogram (XIC) coinciding with the UV peak (Figure 5C). The elution order of antibodies was predicted well by their pI values: eculizumab (pI 6.1), brentuximab-vedotin (predicted pI 6.7), bevacizumab (pI 8.3), trastuzumab (pI 9.1), and NISTmAb (pI 9.2). This order mirrors previous reports. An exception was BSA (pI 4.7), which despite a predicted net negative charge at the starting pH remained bound to the resin and eluted later than eculizumab. Our volatile buffer system also resolved acidic and basic variants of trastuzumab and NISTmAb, generating chromatographic patterns similar to those described in the literature. Notably, the starting low-pH buffer used here was sufficiently acidic to retain eculizumab, which was not observed in prior work.^36,41,42^

**Figure 5.**
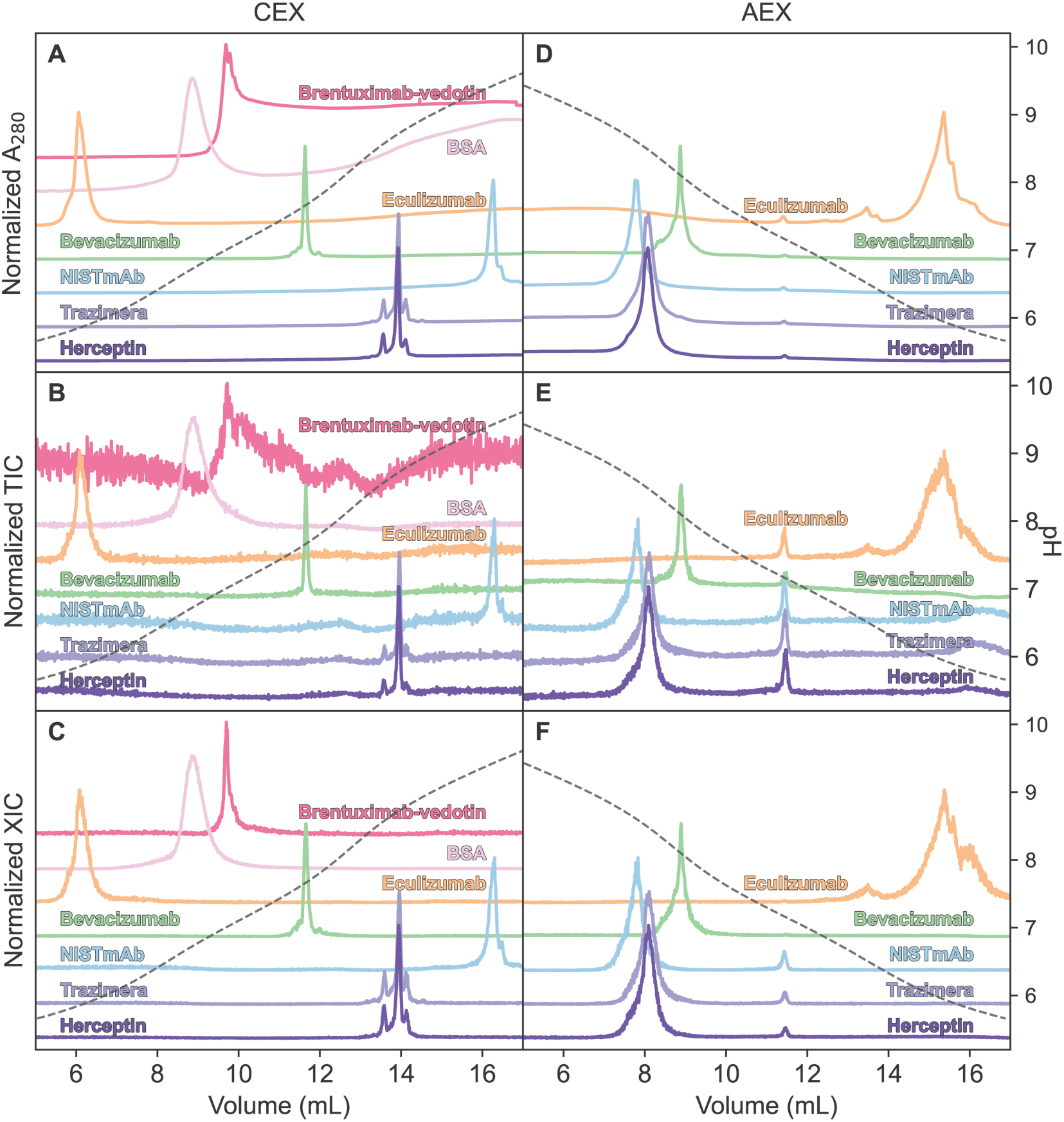
Analysis of volatile compound pH gradients for cation and anion exchange chromatography. The left column of chromatogram plots shows CEX data, measured when a linear gradient from pH 5.46 to pH 9.84 gradient was applied, while the right column of chromatograms shows AEX data recorded by applying a linear gradient form pH 9.84 to pH 5.46. Traces are colour coded for each of the analytes. The top row of plots display the A_280_ measurements made (A, D), while the second row represents the TIC (B, E). The bottom two plots display XICs from the most intense protein mass signals in each spectrum (C, F). Baselines for each analyte have been offset to improve readability of overlapping peaks.

We next applied the same strategy to anion-exchange (AEX) chromatography, inverting the gradient to run from pH 9.84 to 5.46 (Figure 5D-F). As expected, elution order again followed the pI values of the analytes, with NISTmAb eluting first, followed by trastuzumab, bevacizumab, and finally eculizumab. This ranking was consistent with the corresponding CEX chromatograms. Interestingly, in the AEX experiment, eculizumab produced an additional peak and broader chromatographic profile compared to CEX, suggesting subtle differences in protein-resin interactions under the two modes. Spectra obtained throughout both the CEX and AEX pH gradient experiments are of high quality, consistent with previously recorded spectra (Figure S6).

### Hydrophobic Interaction Chromatography (HIC)

We further examined the compatibility of these compounds with HIC. Like IEX, HIC performance is highly sensitive to solution pH, making it a stringent benchmark for evaluating the buffering capabilities of our fluorinated ethylamines. Protein retention on HIC resins is governed by interactions between uncharged surface regions of the protein and the moderately hydrophobic stationary phase. Because the effective net charge of proteins is controlled by buffer pH, precise pH regulation is important in HIC experiments, even if the relationship between retention and pH can be complex.^16^ Traditionally, HIC is performed using phosphate buffers, with elution achieved by an ammonium sulfate gradient to modulate the salting-out effect. However, both phosphate and ammonium sulfate are incompatible with ESI-MS. Previous studies have shown that AmAc can substitute for ammonium sulfate to drive salting-out, though at a high concentration of 4.1 M.^16,43–45^ This precedent motivated us to test the performance of volatile buffer systems in HIC using DFEA as the buffering component and AmAc as the salting-out agent.

Using a Waters Protein-Pak Hi Res HIC column, we observed strong retention of monoclonal antibodies in a mobile phase containing 50 mM DFEA and 4.0 M AmAc at pH 7. Importantly, analyte elution order was preserved between conventional phosphate-based mobile phases (Figure 6A) and the volatile DFEA-AmAc system (Figure 6B). Notably, the bevacizumab peak, which contained an unresolved shoulder in the phosphate buffer, was resolved into two distinct peaks in the volatile mobile phase. Both species had a masses of approximately 149.2 – 149.36 kDa (Figure S7), corresponding to the mixture of G0F and G1F glycoforms of bevacizumab.^46^ Given the absence of significant charge variants or degradation products in our IEX experiments, we attribute this chromatographic separation to conformational heterogeneity of the antibody under HIC conditions.

**Figure 6.**
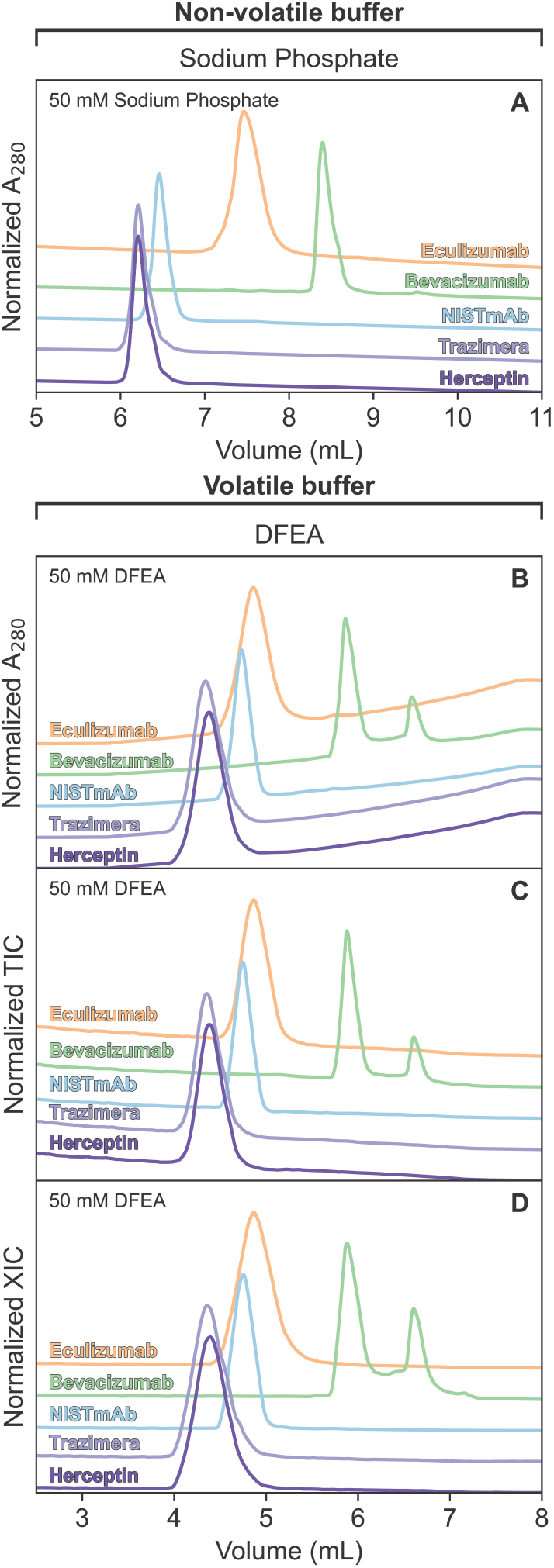
Comparison of HIC for monoclonal antibodies. The topmost plot shows the A_280_ chromatogram (A) obtained using a sodium phosphate – ammonium sulfate-based mobile phase. The next plot represents A_280_ (B), followed by plots displaying TIC (C) and XIC (D), all obtained using a DFEA – ammonium acetate mobile phase. Baselines for each analyte have been offset to improve readability of overlapping peaks.

We further tested brentuximab-vedotin to assess the utility of HIC in volatile buffer systems for DAR analysis. Chromatographic profiles obtained in DFEA-AmAc were highly similar to those from phosphate-based buffers in both peak distributions and relative intensities (Figure 7). Crucially, the use of a volatile mobile phase enabled direct online MS analysis for DAR determination. Whereas IEX-MS spectra of brentuximab-vedotin were often complex, the HIC separation yielded simplified spectra for each peak (Figure S7), indicating that molecular heterogeneity was effectively resolved chromatographically. Deconvolution of these spectra revealed that the earliest-eluting species corresponded to unconjugated brentuximab (148.1 kDa), while subsequent peaks increased in mass by ∼2.6 kDa increments (Figure 7D), consistent with the addition of two monomethylauristatin A molecules (MW = 1316 Da).^47^ Thus, the volatile HIC system enabled direct resolution of four DAR species (DAR-0, DAR-2, DAR-4, and DAR-6), each readily distinguished in the mass spectra (Figure 7D).

**Figure 7.**
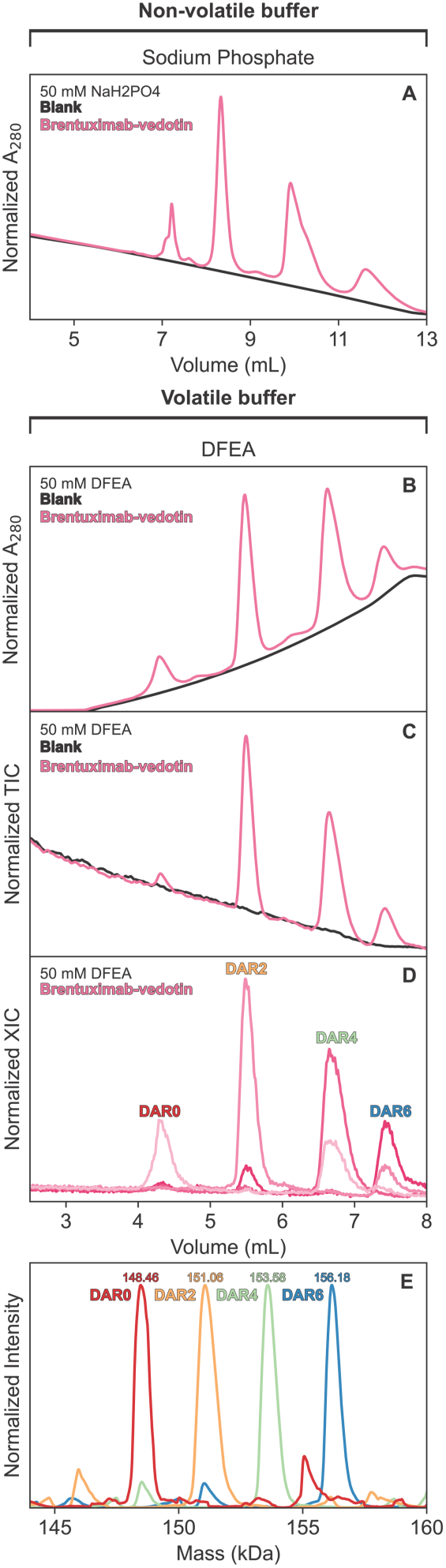
Comparison of HIC for Bretuximab-vedotin examining drug-antibody complexes. The topmost plot (A) shows the A_280_ chromatogram obtained using a sodium phosphate – ammonium sulfate-based mobile phase. The next plot represents A_280_ (B), followed by plots displaying TIC (C) and XIC (D), all obtained using a DFEA mobile phase. The final plot displays the deconvoluted masses obtained from each of the four peaks observed in chromatography, exemplifying the separation of different drug-antibody complexes based on the drug-antibody ratio (DAR).

### Size Exclusion Chromatography (SEC)

Finally, we explored the compatibility of the novel compounds with SEC. SEC in pure ammonium acetate is a well-established LC type in native LC-MS studies of proteins.^24,48,49^ Retention of analytes by size exclusion resin is based on the size and shape of analytes, thus the physical principle of size exclusion is therefore independent of solution pH. Nevertheless, properties of protein analytes, especially macromolecular complexes, are influenced by pH.^50^ For this reason, pH control during a SEC run is still advantageous. We sought to demonstrate compatibility of our solvent system with the SEC-MS workflow. We analyzed a set of test proteins: cytochrome *c*, BSA, and our five antibodies (NISTmAb, eculizumab, bevacizumab, and two trastuzumab samples from different manufacturers) and one antibody-drug conjugate (brentuximab-vedotin) on ACQUITY Premier Protein SEC 250Å column (Waters) and recorded ESI-MS TICs in a fully volatile buffer containing 20 mM DFEA (pH 7.0) and 200 mM ammonium acetate. The chromatograms were very similar to those measured when the same analytes were analyzed in a traditional non-volatile buffer system containing 20 mM HEPES (pH 7.0) with 200 mM NaCl (Figure 8). Elution rank of proteins matched our expectations based on molecular weights: 12 kDa Cytochrome *c* eluted last, preceded by the 67 kDa BSA monomer, and finally preceded by the ∼150 kDa mAbs. Obtained spectra were of high quality, and consistent with spectra obtained using other chromatography methods (Figure S8).

**Figure 8.**
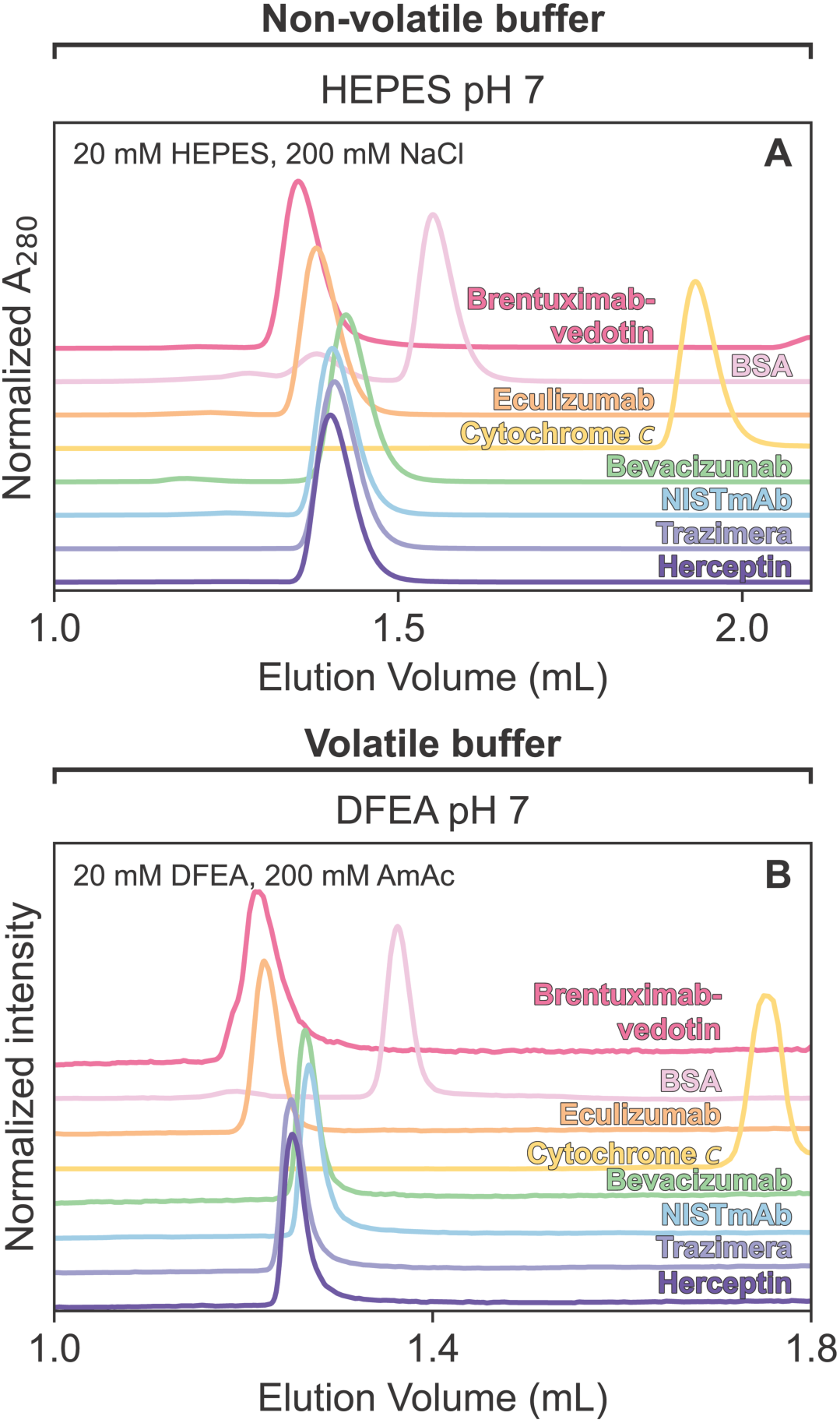
Comparison of SEC in volatile and non-volatile mobile phases at pH 7. Traces are coloured for each of the analytes. Panel A demonstrates the chromatogram obtained using non-volatile components HEPES and NaCl, while panel B is the chromatogram using the same parameters but obtained with a volatile mobile phase containing DFEA and AmAc.

## Discussion and Conclusions

Here we demonstrated that mobile phases composed of MFEA, DFEA, TFEA, and AmAc are compatible with the analysis of mAbs, ADCs, and other proteins such as BSA and cytochrome *c* as part of CEX, AEX, HIC, and SEC workflows. As expected, steep gradients allowed for rapid method scouting, while shallow gradients provided higher resolution separations. When comparing the volatile mobile phases with traditional non-volatile buffers, we observed only minor differences, such as slight shifts in retention times and modest peak broadening. In most cases, analytes eluted earlier in the volatile systems. Importantly, coupling these volatile systems to online ESI-MS enabled direct measurement of mass spectra for mAbs and ADCs under fully native conditions (Figures 2, and 4; Figures S2, S4, S6-S8). We note that volatile and non-volatile experiments were performed on different LC instruments with non-identical dwell and dead volumes, contributing to apparent offsets in retention times. Because these offsets have no bearing on the utility of the proposed mobile phases, we did not attempt to correct for them.

A central application of analytical workflows in the development of biotherapeutics is the demonstration of biosimilarity between innovator and biosimilar products.^18,19,51–55^ To assess the applicability of the volatile mobile phases for this purpose, we included both innovator trastuzumab (Herceptin) and its biosimilar (Trazimera) in our analyses. In both volatile and non-volatile buffers, chromatographic behavior, including retention times, peak shapes, and the appearance of minor variants, were identical between the two products. Mass spectrometry further confirmed equivalent molecular weights and indistinguishable spectral fine structures, including glycosylation heterogeneity. We also established volatile buffer formulations for pH-gradient elution, maintaining consistently low ionic strength (<40 mM) throughout the gradient. These systems proved effective for both CEX and AEX chromatography. The elution order of mAbs was predictable and consistent with literature precedents, with one notable exception: BSA eluted after eculizumab in CEX, despite its lower pI and predicted negative charge at the starting pH. Such unexpected behaviors highlight the value of combining CEX and AEX analyses, as the two modes provide complementary information. For example, while acidic mAbs such as eculizumab were weakly retained under CEX conditions and eluted without resolving charge variants, basic mAbs such as trastuzumab and NISTmAb showed clear variant resolution. In AEX, the scenario was inverted with basic mAbs eluted early, while eculizumab remained bound and displayed distinct charge variant peaks. Crucially, the MS-compatible volatile mobile phases allowed straightforward mass determination of all resolved analytes. These results underscore the potential of volatile buffer systems to support biosimilarity assessments by providing a platform that integrates chromatographic resolution with detailed MS-based molecular characterization.

We demonstrate mono-, di-, and tri-fluoroethylamines as promising MS-compatible buffering agents for LC-MS applications. In combination with AmAc, these compounds can provide effective buffering across a broad pH range of 3.6-10.3, thereby enabling LC-MS workflows for the study of pH-sensitive proteins, antibody quality control, and ligand interactions. While our results demonstrate that volatile mobile phases largely reproduce chromatographic trends observed with conventional buffers, minor shifts in retention order and peak behavior warrant further study to ensure analyte conformation and charge are preserved. One current limitation is the lack of readily available MS-grade reagents for fluorinated ethylamines, which can introduce adducts or contaminant peaks. Despite this, the ability of these volatile systems to maintain resolution of distinct DAR species in brentuximab-vedotin highlights their strong potential for advancing native LC-MS. In summary, we establish fluorinated ethylamines as a universal class of volatile buffering agents that extend the reach of native LC-MS, bridging longstanding limitations of AmAc and opening new possibilities for high-resolution analysis of complex, pH-sensitive biomolecules.

## Acknowledgements

K.R.V., and B.T.V.D. acknowledge support from a Graduate Tuition Scholarship from the University of Guelph. MS data were recorded at the Mass Spectrometry Facility of the Advanced Analysis Centre, University of Guelph. We thank Dr. Dyanne Brewer (University of Guelph) for assistance with MS measurements. Financial support was provided by a Discovery Grant from the Natural Sciences and Engineering Research Council (NSERC) of Canada (RGPIN-2021-02843) to S.V, a NSERC Discovery Grant (RGPIN 2020−05043) to D.K.O., a NSERC Alliance Catalyst Grant (ALLRP 57555822) to D.K.O., and Canadian Foundation for Innovation - John R. Evans Leaders Fund (Project 43757) to D.K.O. J.S.S was supported by a NSERC Canada Graduate Research Scholarship – Master’s program (CGS-M).

## Materials and Methods

### Materials

Ethylamine aqueous solution (471208-100ML) was obtained from Millipore Sigma; 2-fluoroethylamine (MFEA) hydrochloride (A155303) from Ambeed, 2,2-difluoroethylamine (DFEA) (SC-43926) from StruChem (Wujiang, China); 2,2,2-trifluoroethylamine (TFEA) (AC303500100), LC-MS acetic acid (A113), LC-MS water (W6-4), and LC-MS ammonium acetate (A114-50) were from Fisher Scientific. Bovine heart cytochrome c (C3131) was from Millipore Sigma, BSA (800-095-EG) from Wisent, and NISTmAb (8671) from NIST. Aliquots of clinical samples of brentuximab-vedotin (Adcetris), eculizumab (Soliris), bevacizumab (Avastin), innovator trastuzumab (Herceptin), and biosimilar trastuzumab (Trazimera) were purchased from Evidentic GmbH (Berlin, Germany). Protein-Pak Hi Res SP (100 × 4.6 mm, 7 µm particle size, P/N 186004930), Protein-Pak Hi Res Q (100 × 4.6 mm, 5 µm particle size, P/N 186004931), Protein-Pak Hi Res HIC (100 × 4.6 mm, 2.5 µm particle size, P/N 186007583), and ACQUITY UPLC Protein BEH SEC, 200 Å (150 × 4.6 mm, 1.7 µm particle size, P/N 186005225) columns were from Waters.

Hydrochloride-free MFEA was produced by stirring a solution of 2-fluoroethanammonium chloride (5.0 g, 50 mmol) in purified water (100 mL) to which sodium hydroxide (30 g, 750 mmol) was added slowly (over 20 min) at 0 °C. The solution was allowed to stir for 20 min at 0 °C. The desired product was vacuum distilled into a trap residing in liquid nitrogen and used without further purification. This was characterized by ^1^H and ^19^F 1D NMR as well as direct infusion MS to ensure purity, showing the product was free of HCl but not free of water. For the purposes of generating aqueous solutions this was not a concern.

For IEX experiments, 1 mg/mL samples of each analyte were prepared by diluting antibody or ADC aliquots in 20 mM DFEA, pH 7.0 buffer, or by dissolving solid powder of BSA and cytochrome *c* in the same buffer. For HIC, samples of antibodies and the ADC were diluted in 50:50 mixture of mobile phases A and B to 2.5 mg/mL final concentration. 10 µL of analyte sample were injected per analysis. Mobile phases for salt gradient-based ion exchange chromatography contained 20 mM buffering compound in buffer A and B at a desired pH, and 1 M salt in buffer B only. HIC mobile phases contained 50 mM buffering compound at pH 7.0 in buffer A and B, and 1.2 M ammonium sulfate or 4 M ammonium acetate in buffer A only. Final pH was adjusted with concentrated HCl or NaOH in non-volatile mobile phases. Volatile mobile phases were prepared by dissolving free amine form of MFEA/DFEA/TFEA and ammonium acetate as required, pH was then adjusted with LC-MS grade AcOH or NH_4_OH. For ion exchange pH gradient elution, a low pH buffer was prepared by mixing 40 mM AcOH, 20 mM TFEA, 20 mM DFEA, with a final pH of 5.46. High pH buffer contained 32 mM AcOH, 20 mM DFEA, 20 mM MFEA, and 30 mM ethylamine, with a final pH of 9.84. For SEC mobile phases, 20 mM of DFEA was used when making pH 7 solutions, 200 mM of AmAc added to all buffers, with pH adjusted as previously described. Exact composition of mobile phases for all chromatography types is listed in Table 1.

**Table 1:**
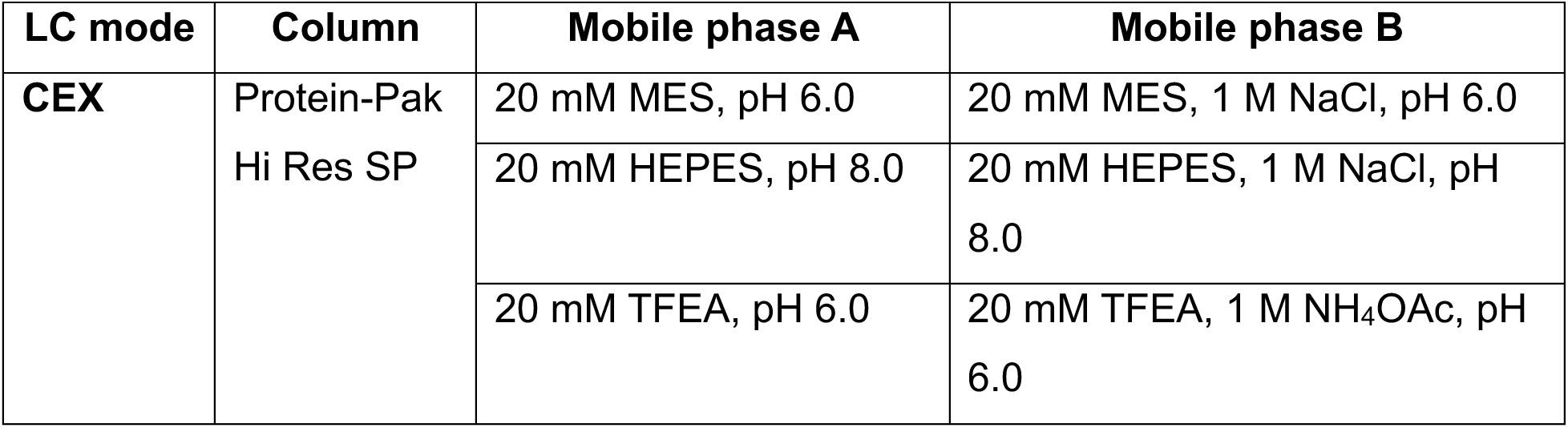
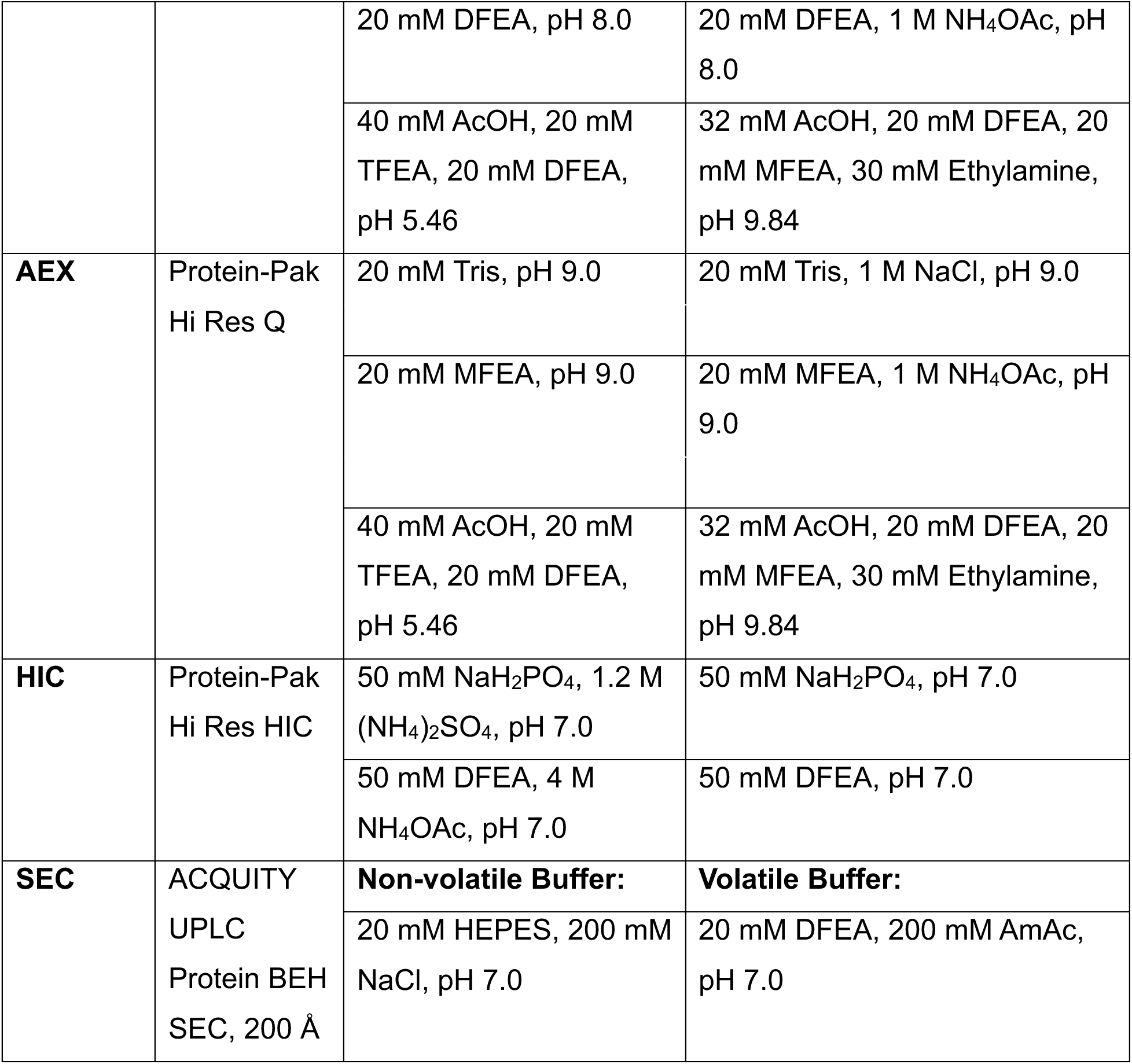
Buffering solutions used in each chromatography methodology.

All MS measurements were performed in positive-ion mode on a Waters Synapt G2Si instrument with a standard ESI source. Spectra were recorded over an 800-8000 *m/z* range, on a time-of-flight mass analyzer set to resolution mode. The MS instrument was externally calibrated using the ESI-L tune mix (G1969-8500) from Agilent. Exact LC settings used are available in Tables S1-S5.

### pH titrations

A potentiometric titration of MFEA was performed to validate the theoretical p*K*_a_ value of 9.0, conducted with a Fisher Scientific Accument AB15 pH meter at room temperature. Standard curve and pH titration measurements were obtained in mV mode, and 5 pH standards (2, 4, 7, 10, and 12) were used to produce the standard curve for the conversion of mV values to pH. For the pH titration of MFEA the initial volume was 20 mL with a concentration of 100 mM while NaOH at 0.2N was used as a titrant. p*K*_a_ values were then determined by fitting the potentiometric titration data to an analytical equation accounting for a single p*K*_a_ value using scripts written in house in Python 3.8 using the lmfit package.^24^ To determine uncertainty in the value of the fitted p*K*_a_ of MFEA we performed a Monte Carlo analysis, introducing random errors, determined by the root-mean-square deviation between the experimentally observed points and those derived from the optimal-fit model, into the best-fit model. Obtained values were then represented in a histogram, which was then fit to a normal distribution to calculate the mean expectation value and standard deviation. We present the uncertainties as 2 standard deviations in the derived values, representing a 95% confidence interval. All relevant scripts are available upon request.

