## Supporting Information for "A universal buffer system for native LC-MS analysis of antibody-based therapeutics"

**This file contains:**

**Tables S1-5:** Detailed chromatography parameters

**Figure S1:** Cation exchange chromatography at pH 6 and 8 with a steep ion gradient

**Figure S2:** Additional CEX shallow salt gradient mass spectra

**Figure S3:** Potentiometric titration of MFEA

**Figure S4:** Additional AEX shallow salt gradient mass spectra

**Figure S5:** Anion exchange chromatography at pH 9 with a steep ion gradient

**Figure S6:** Mass spectra recorded during CEX and AEX with a pH gradient

**Figure S7:** Mass spectra recorded during HIC

**Figure S8:** Mass spectra recorded during SEC

**Table S1:** Cation exchange chromatography parameters

| LC Experiment: Analyte | Flow Rate (mL / min) | Time (min) | % Mobile Phase A | % Mobile Phase B |
|---|---|---|---|---|
| **Shallow gradient:** Cytochrome *c* (pH 8)<br>Bevacizumab, NISTmAb, Trazimera, Herceptin (pH 6 and 8) | 0.6 | 0 | 100 | 0 |
| | | 6 | 100 | 0 |
| | | 26 | 80 | 20 |
| | | 26.01 | 0 | 100 |
| | | 30 | 0 | 100 |
| | | 30.01 | 100 | 0 |
| | | 38 | 100 | 0 |
| **Shallow gradient:** Cytochrome C (pH 6) | 0.6 | 0 | 100 | 0 |
| | | 2 | 100 | 0 |
| | | 2.01 | 90 | 10 |
| | | 6 | 90 | 10 |
| | | 26 | 70 | 30 |
| | | 26.01 | 0 | 100 |
| | | 30 | 0 | 100 |
| | | 30.01 | 100 | 0 |
| | | 38 | 100 | 0 |
| **Steep gradient:** Cytochrome C, Bevacizumab, NISTmAb, Trazimera, Herceptin | 0.6 | 0 | 100 | 0 |
| | | 6 | 100 | 0 |
| | | 20 | 0 | 100 |
| | | 24 | 0 | 100 |
| | | 24.01 | 100 | 0 |
| | | 32 | 100 | 0 |

**Table S2:** Anion exchange chromatography parameters

| LC Experiment: Analyte | Flow Rate (mL / min) | Time (min) | % Mobile Phase A | % Mobile Phase B |
|---|---|---|---|---|
| **Shallow gradient:** Brentuximab-vedotin, BSA (pH 9) | 0.4 | 0 | 100 | 0 |
| | | 1 | 100 | 0 |
| | | 1.01 | 80 | 20 |
| | | 8 | 80 | 20 |
| | | 38 | 60 | 40 |
| | | 38.01 | 0 | 100 |
| | | 42 | 0 | 100 |
| | | 42.01 | 100 | 0 |
| | | 50 | 100 | 0 |
| **Shallow gradient:** Eculizumab (pH 7) Bevacizumab, NISTmAb, Trazimera, Herceptin (pH 7 and 9) | 0.4 | 0 | 100 | 0 |
| | | 8 | 100 | 0 |
| | | 38 | 80 | 20 |
| | | 38.01 | 0 | 100 |
| | | 42 | 0 | 100 |
| | | 42.01 | 100 | 0 |
| | | 50 | 100 | 0 |
| **Shallow gradient:** Eculizumab (pH 9) | 0.4 | 0 | 100 | 0 |
| | | 1 | 100 | 0 |
| | | 1.01 | 90 | 10 |
| | | 8 | 90 | 10 |
| | | 38 | 70 | 30 |
| | | 38.01 | 0 | 100 |
| | | 42 | 0 | 100 |
| | | 42.01 | 100 | 0 |
| | | 50 | 100 | 0 |
| **Shallow gradient:** Brentuximab-vedotin, BSA (pH 7) | 0.4 | 0 | 100 | 0 |
| | | 1 | 100 | 0 |

| | | | | |
|---|---|---|---|---|
| | | 1.01 | 95 | 5 |
| | | 8 | 95 | 5 |
| | | 38 | 75 | 25 |
| | | 38.01 | 0 | 100 |
| | | 42 | 0 | 100 |
| | | 42.01 | 100 | 0 |
| | | 50 | 100 | 0 |
| **Steep gradient:** Brentuximab-vedotin, BSA, Eculizumab, Bevacizumab, NISTmAb, Trazimera, Herceptin | 0.4 | 0 | 100 | 0 |
| | | 6 | 100 | 0 |
| | | 20 | 0 | 100 |
| | | 24 | 0 | 100 |
| | | 24.01 | 100 | 0 |
| | | 36 | 100 | 0 |

**Table S3:** pH gradient IEX parameters

| LC Experiment: Analyte | Flow Rate (mL / min) | Time (min) | % Mobile Phase A | % Mobile Phase B |
|---|---|---|---|---|
| **CEX pH gradient:** Brentuximab-vedotin, BSA, Eculizumab, Bevacizumab, NISTmAb, Trazimera, Herceptin | 0.6 | 0 | 100 | 0 |
| | | 6 | 100 | 0 |
| | | 26 | 0 | 100 |
| | | 30 | 0 | 100 |
| | | 30.01 | 100 | 0 |
| | | 38 | 100 | 0 |
| **AEX pH gradient:** Eculizumab, Bevacizumab, NISTmAb, Trazimera, Herceptin | 0.4 | 0 | 0 | 100 |
| | | 8 | 0 | 100 |
| | | 38 | 100 | 0 |
| | | 46 | 100 | 0 |
| | | 46.01 | 0 | 100 |
| | | 58 | 0 | 100 |

**Table S4:** Hydrophobic interaction chromatography parameters

| Analyte | Flow Rate (mL / min) | Time (min) | % Mobile Phase A | % Mobile Phase B |
|---|---|---|---|---|
| Eculizumab, Bevacizumab, NISTmAb, Trazimera, Herceptin, Brentuximab-vedotin | 0.3 | 0 | 100 | 0 |
| | | 8 | 100 | 0 |
| | | 23 | 0 | 100 |
| | | 31 | 0 | 100 |
| | | 31.01 | 100 | 0 |
| | | 40 | 100 | 0 |

**Table S5:** Size exclusion chromatography parameters

| Analyte | Flow Rate (mL / min) | Time (min) | % Mobile Phase A | % Mobile Phase B |
|---|---|---|---|---|
| Cytochrome C (pH 7) | 0.3 | 0 | 100 | 0 |
| | | 10 | 100 | 0 |
| Brentuximab-vedotin, BSA, Eculizumab, Bevacizumab, NISTmAb, Trazimera, Herceptin (pH 7) | 0.03 | 0 | 100 | 0 |
| | | 62 | 100 | 0 |
| | 0.3 | 62.2 | 100 | 0 |
| | | 66 | 100 | 0 |

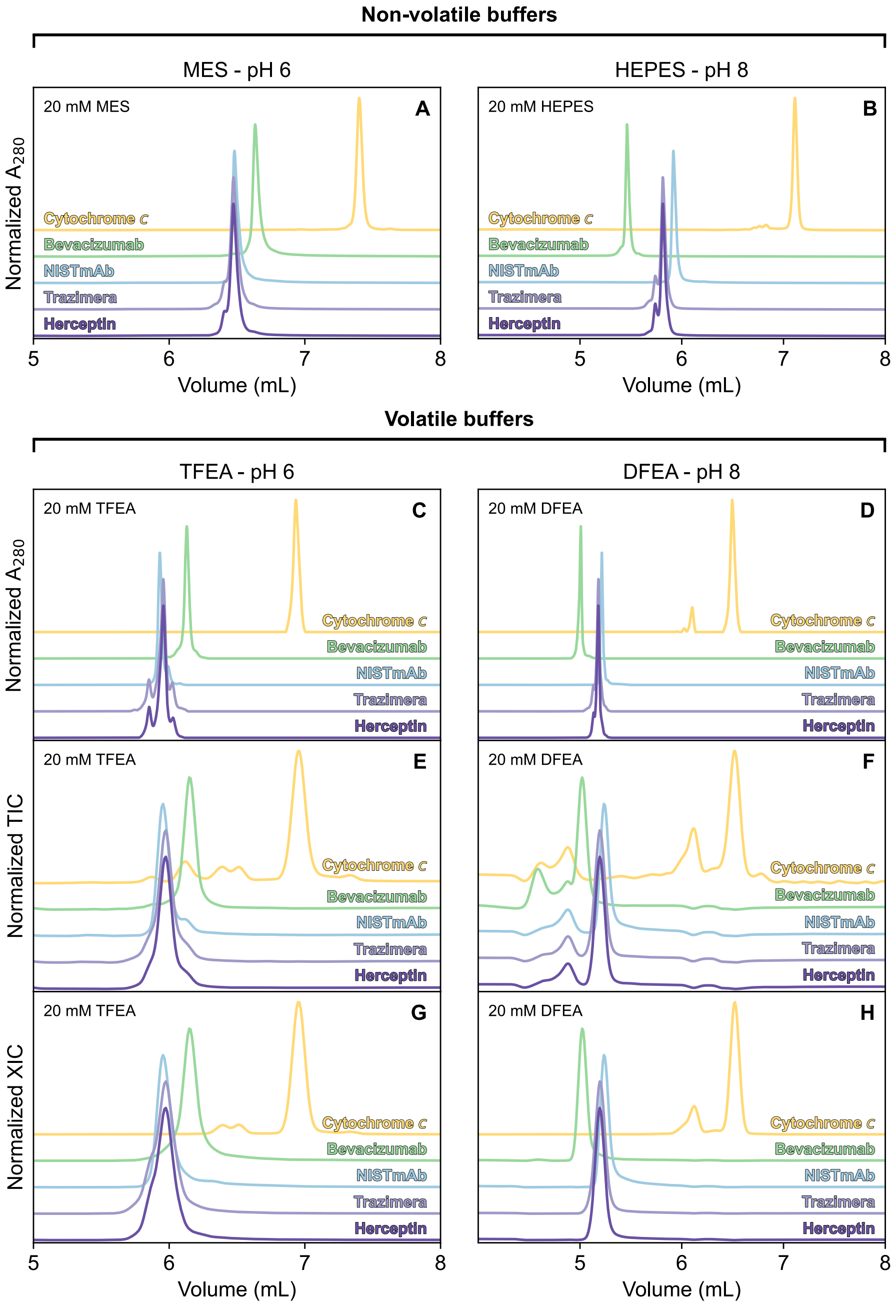


**Figure S1. Cation exchange chromatography at pH 6 and 8 with a steep ion gradient.** The first row of plots displays $A_{280}$ chromatograms obtained with an MES mobile phase (pH 6.0) or a HEPES mobile phase (pH 8.0), and a steep NaCl gradient. The next rows of plots show $A_{280}$, TIC, and XIC chromatograms obtained with either a TFEA mobile phase (pH 6.0) or a DFEA mobile phase (pH 8.0), also using a steep ionic strength gradient produced with AmAc. Baselines for each analyte have been offset to improve readability of overlapping peaks. Traces are colour coded for each analyte.

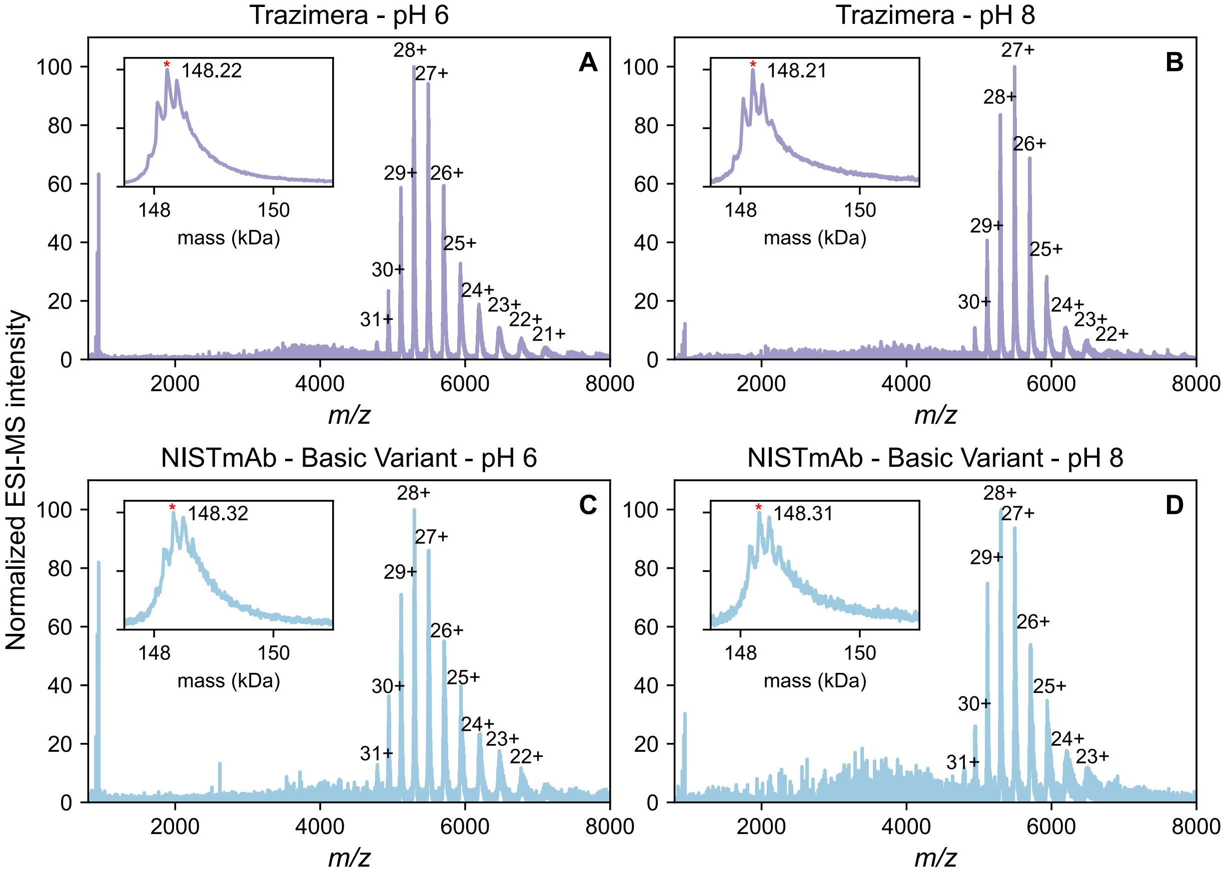


**Figure S2. Additional CEX shallow salt gradient mass spectra.** Traces are colour coded for each analyte. Prominent charge states are labelled and deconvoluted masses are displayed as inset plots. The NISTmAb spectra represents the secondary peak seen in the pH 6 and pH 8 CEX chromatograms, accounting for a basic variant with a mass difference of approximately 130 Da.

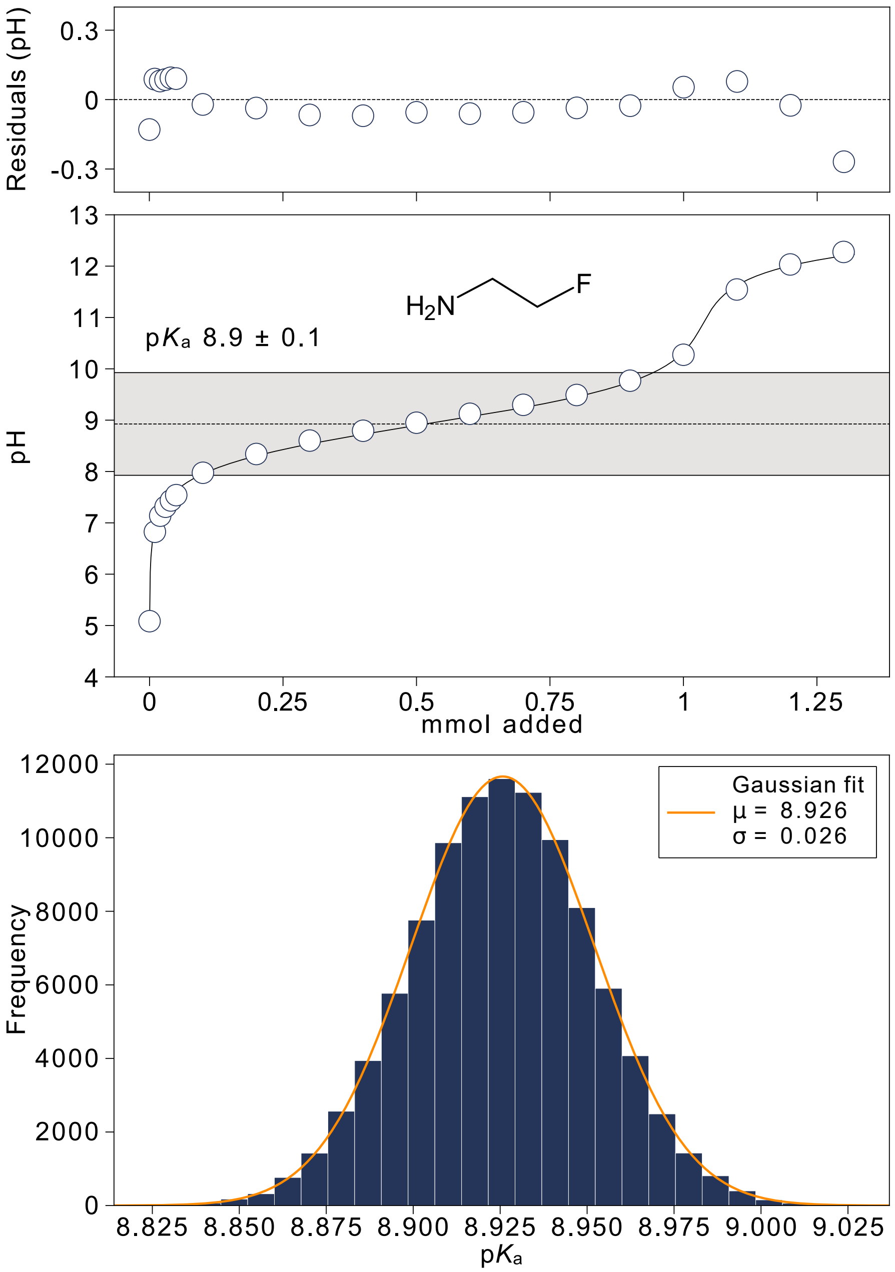


**Figure S3. Potentiometric titration of MFEA.** pH values were measured in a solution of MFEA hydrochloride titrated with sodium hydroxide and were fit to a mathematical model described previously.[24] Solid line is the plot of model function using the best fit parameters. Gray region indicates the interval where MFEA provides effective pH control, based on ±1 pH unit from the fitted p$K_a$ value. Fitting residuals are shown in the top panel, and the bottom panel displays a Monte Carlo histogram depicting the uncertainty in the value of fitted p$K_a$.

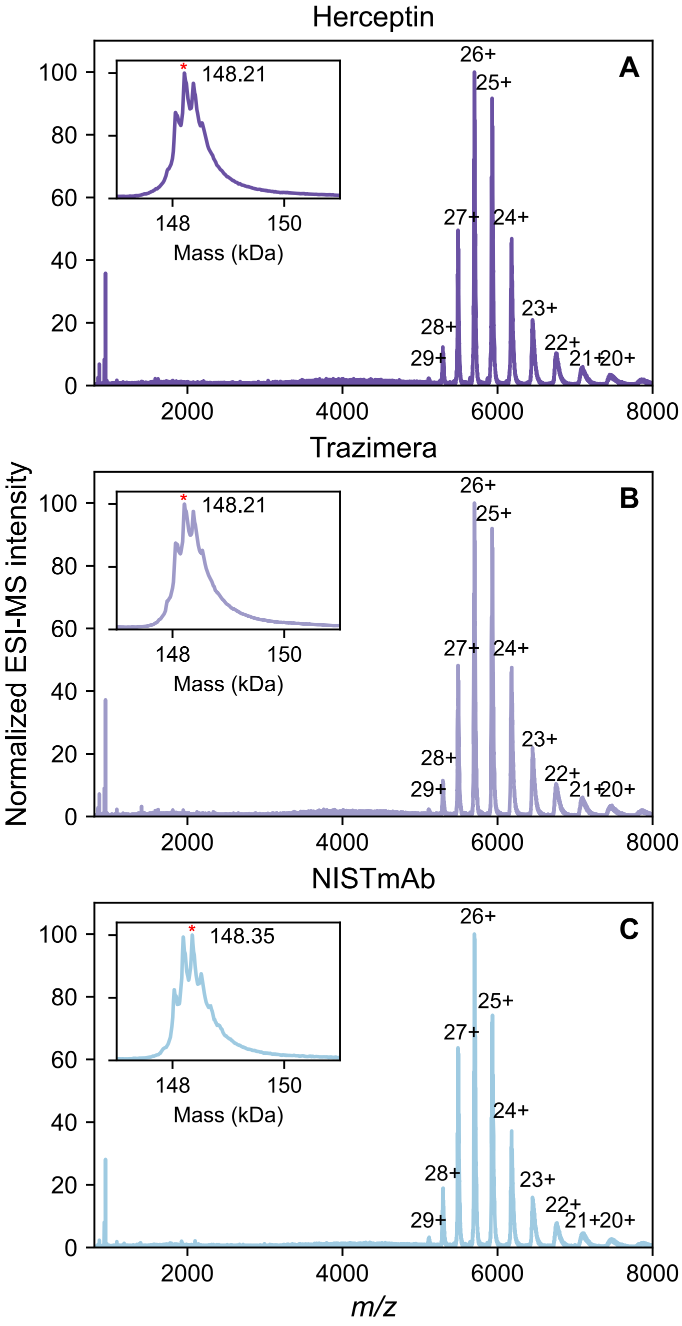


**Figure S4. Additional AEX shallow salt gradient mass spectra.** Prominent charge states are labelled and deconvoluted masses are displayed as inset plots. Spectral and deconvoluted mass traces are colour coded for each analyte.

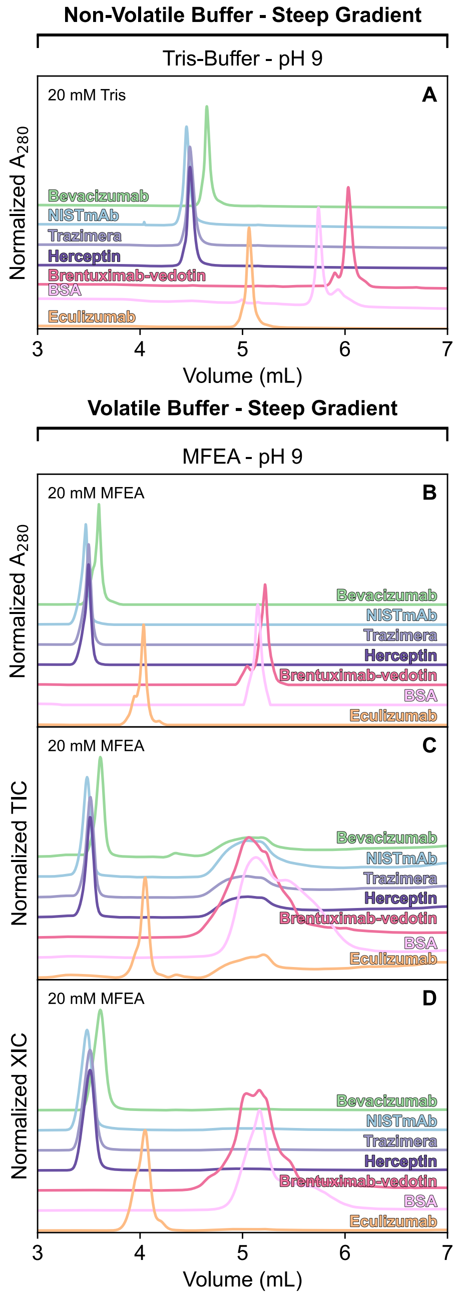


**Figure S5. Anion exchange chromatography at pH 9 with a steep ion gradient.** Trastuzumab is coloured purple, NISTmAb is shown in blue, bevacizumab in green, eculizumab in orange, BSA in light pink, and brentuximab-vedotin in dark pink. The first plot displays an $A_{280}$ chromatogram obtained in a non-volatile mobile phase, Tris at pH 9, with a steep ionic strength gradient. The $A_{280}$, TIC, and XIC chromatograms obtained in MFEA buffer at pH 9 are shown in the subsequent plots. Baselines for each analyte have been offset to improve readability of overlapping peaks.

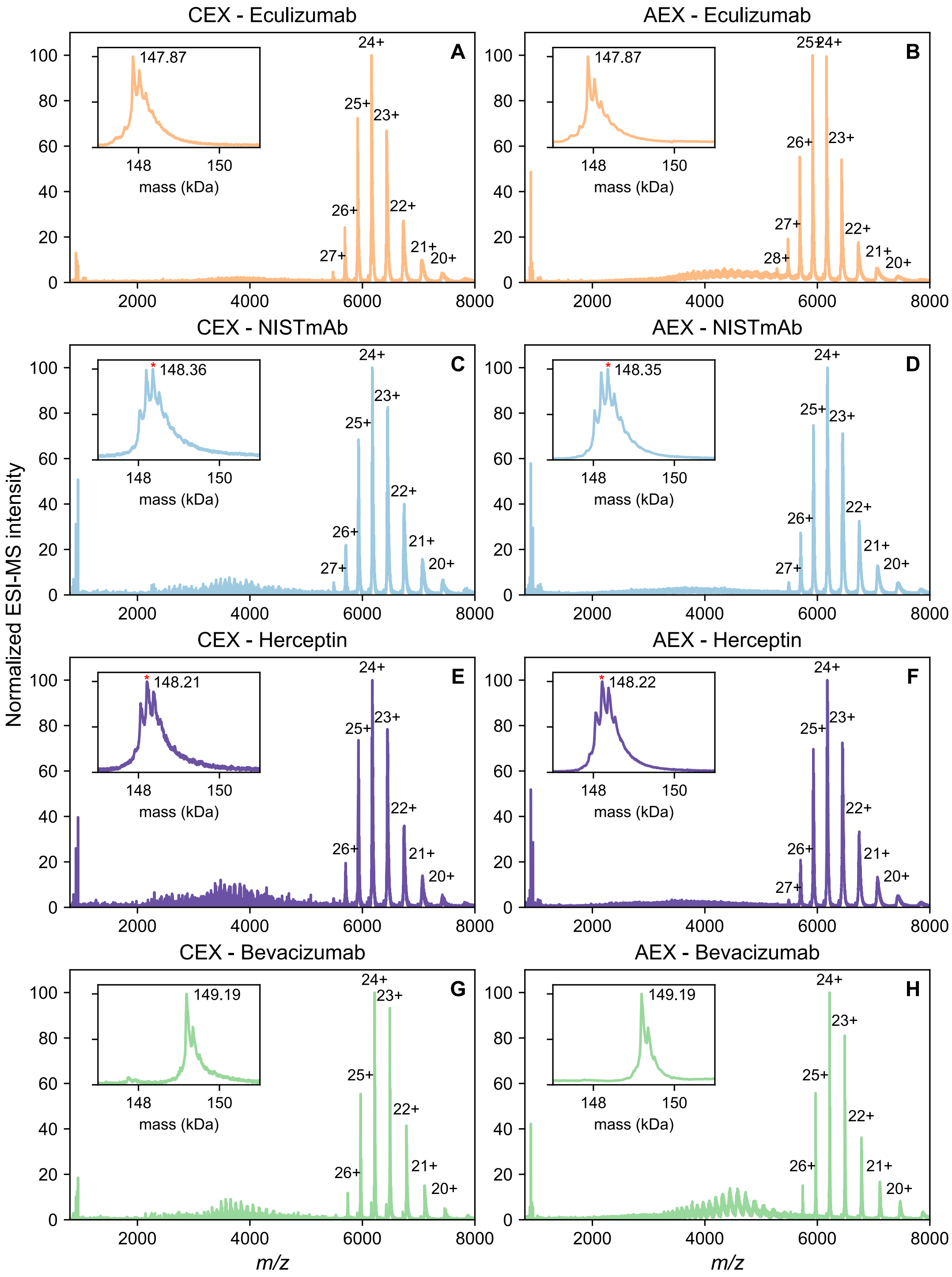


**Figure S6. Mass spectra recorded during CEX and AEX with a pH gradient.** Eculizumab is coloured orange, NISTmAb is coloured blue, trastuzumab from Herceptin is coloured purple, and bevacizumab is coloured green. Prominent charge states are labelled and deconvoluted masses are displayed as inset plots.

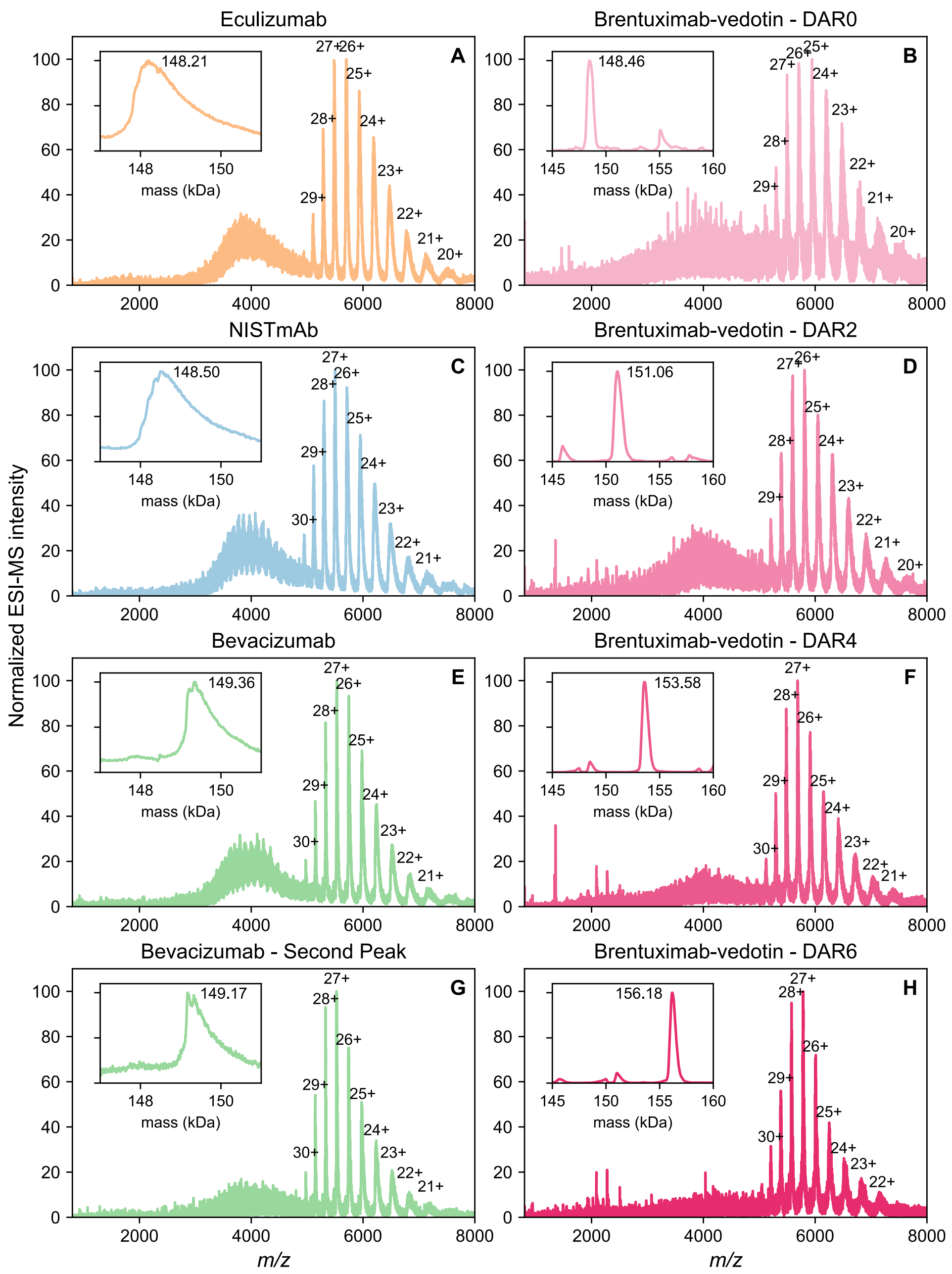


**Figure S7. Mass spectra recorded during HIC.** Eculizumab is coloured orange, NISTmAb is coloured blue, and bevacizumab is coloured green. Brentuximab-vedotin DAR variants are coloured a gradient of dark to light pink to align with the deconvoluted masses displayed in Figure 7. Prominent charge states are labelled and deconvoluted masses are displayed as inset plots. Spectra from both peaks of Bevacizumab are displayed, demonstrating both have similar charge states and nearly identical deconvoluted masses.

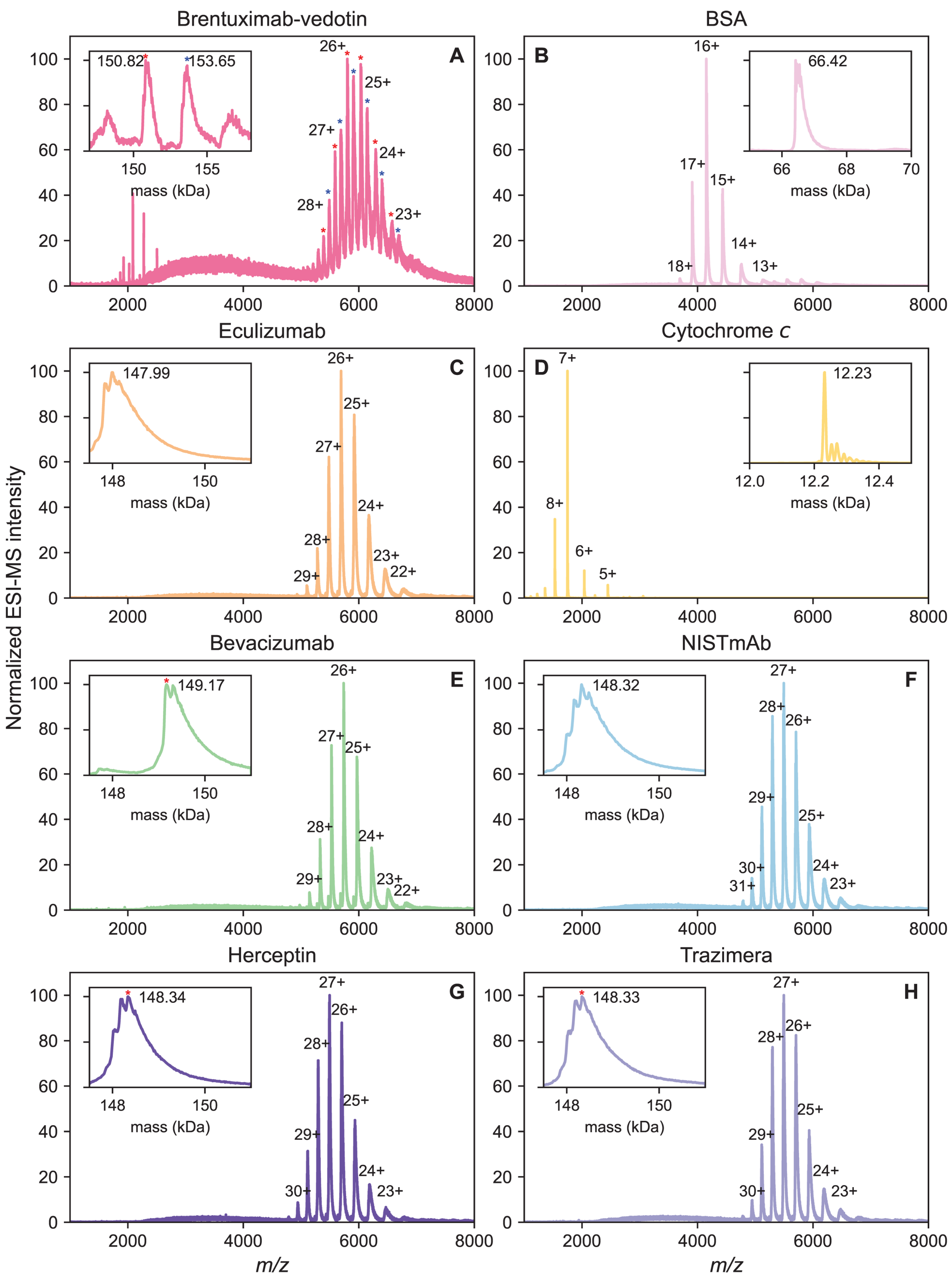


**Figure S8. Mass spectra recorded during SEC.** Brentuximab-vedotin is coloured dark pink, BSA is coloured pink, eculizumab is coloured orange, cytochrome c is coloured yellow, bevacizumab is coloured green, NISTmAb is coloured blue, and trastuzumab from Herceptin and Trazimera are coloured shades of purple. Prominent charge states are labelled and deconvoluted masses are displayed as inset plots.
